# Diatom – lipid – copepod nexus under threat by global change

**DOI:** 10.1101/2025.08.22.667900

**Authors:** Sigrún Huld Jónasdóttir, Richard Broughton, Andre W. Visser

## Abstract

The central role of lipids in the ecology of high latitude seas is under threat by global change. This hinges on tight ecological links between diatoms and copepods of the genus *Calanus*, that produce and accumulate energy rich lipids. Strongly coupled through seasonal water column stability, nutrient supply, spring bloom succession and overwintering life strategies, this nexus is foundational to the highly productive food webs of Arctic/sub-Arctic North Atlantic. We report that the ability of *Calanus finmarchicus* to accumulate lipids is seriously impaired by changing plankton community and by rising temperatures, being 3 times higher on diatoms than flagellate based diets while suffering around a 60% reduction associated with a 2oC temperature change. This together with higher respiration rates, smaller size at maturity and reduced diatom productivity suggests their ability to overwinter will be compromised, threatening not only their contribution to regional productivity but also in maintaining a significant pool of sequestered carbon via the lipid pump. Similar lipid mediated couplings are prevalent throughout high latitude seas, bringing uncertainty to the ecosystem services they provide.

## Main Text

Global change is reducing the dominance of diatoms in seasonal productivity cycles (1–3). Freshening of surface polar and sub-polar seas is evident as a consequence of climate driven glacial and sea-ice melt (1) and surface warming results in increased stratification reducing the annual replenishment of nutrients from depth to the sunlit surface ocean. This pushes new production to greater depths, making surface productivity more dependent on recycled nutrients and shifts primary producers away from diatoms and towards smaller phytoplankton cells (3). Depending on the depth of the stratified layer, a deep chlorophyll maximum may become light limited, directing the production towards hetero- and mixotrophic phytoplankton, away from autotrophic diatoms, significantly impacting food web transfer efficiency. Importantly, since diatoms are a rich source of lipids (typically 25% of carbon mass (4)), the availability of lipids to these polar and sub-polar ecosystems will be reduced.

Lipid rich copepods are the foundation of high latitude marine ecosystems and form a direct link between primary production by diatoms and other phytoplankton to fish, seabirds and marine mammals (5, 6). Members of the genus *Calanus* are particularly conspicuous due to their large size (2-7 mm), high abundance and energy content. Their success, reflected in their high abundance, stems from their specific life history trait to accumulate lipids (wax esters) and enter diapause (hibernation) at depth for up to 9 months of the year, avoiding high predation risk at the surface when food availability is limited (7). This life history trait is commonly adopted by copepods in highly seasonal environments such as polar ecosystems, monsoons, and upwelling areas (8) and drives the lipid pump (9). The accumulation of lipids is crucial, both as an energy source but also to maintain neutral buoyancy while in a torpid state (10). In high latitude regions *Calanus finmarchicus* starts lipid accumulation after metamorphosis from nauplii to copepodite and reaches maximum level in copepod stage 5 (C5) after which they descend for diapause. The amount and type of lipids as well as overwintering temperatures determine the length of diapause (11), and the faster they can build the requisite lipid reserves, the sooner they can descend for safety at depth (12).

## Results and Discussion

We tested lipid accumulation rates of *C. finmarchicus* reared from stage C1 to C5 on pure diets. Our interest was in quantifying animal size and lipid mass, rather than lipid ratios as is usual in these types of studies. The diets represented the predicted change from a diatom to dinoflagellate and flagellate-based food web. The trials also tested temperature change of 2 degrees from 5 to 7°C. The results show clearly that diatoms are crucial to the lipid accumulation strategy of polar copepods. Stages C4 and C5 were significantly smaller (by 19 and 14% respectively) when grown at 7°C compared to 5°C (Figure 1a). Copepods feeding on the diatom *Thalassiosira weissflogii* (Tw) had notably larger lipid sacks compared to those feeding on the cryptophyte *Rhodomonas salina* (Rs) and the dinoflagellate *Heterocapsa triquetra* (Ht, Fig.2). When translated to wax esters, diatom fed C4 and C5s had 3–15 times higher WE content and filled up to 66% of their size-based capacity compared to a maximum of 12% for those fed on *Rs* and *Ht* (Fig.1b, Table 1). However, the saturation and carbon indexes of the copepod WE reserve did not differ between the treatments (Fig 1c). Temperature did not affect lipid sac size of the C5s but was significantly lower at 7°C compared to 5°C for C4s.

**Table 1.**
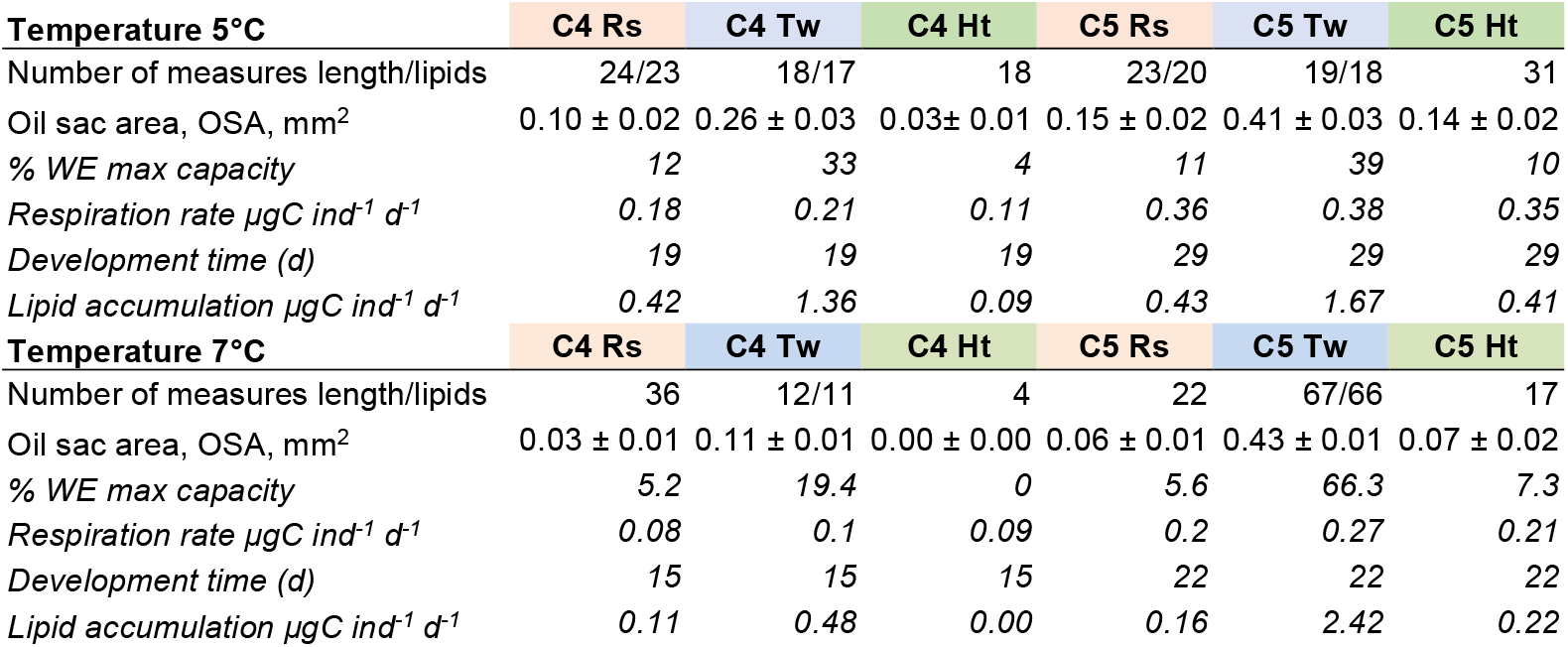
Summary of results for *Calanus finmarchicus* stages C4 and C5 at 7 and 5°C feeding trials. Calculations in italics explained in SI.

| **Temperature 5°C** | **C4 Rs** | **C4 Tw** | **C4 Ht** | **C5 Rs** | **C5 Tw** | **C5 Ht** |
| --- | --- | --- | --- | --- | --- | --- |
| Number of measures length/lipids | 24/23 | 18/17 | 18 | 23/20 | 19/18 | 31 |
| Oil sac area, OSA, mm<sup>2</sup> | 0.10 ± 0.02 | 0.26 ± 0.03 | 0.03 ± 0.01 | 0.15 ± 0.02 | 0.41 ± 0.03 | 0.14 ± 0.02 |
| % WE max capacity | 12 | 33 | 4 | 11 | 39 | 10 |
| Respiration rate $\mu\text{gC ind}^{-1} \text{d}^{-1}$ | 0.18 | 0.21 | 0.11 | 0.36 | 0.38 | 0.35 |
| Development time (d) | 19 | 19 | 19 | 29 | 29 | 29 |
| Lipid accumulation $\mu\text{gC ind}^{-1} \text{d}^{-1}$ | 0.42 | 1.36 | 0.09 | 0.43 | 1.67 | 0.41 |
| **Temperature 7°C** | **C4 Rs** | **C4 Tw** | **C4 Ht** | **C5 Rs** | **C5 Tw** | **C5 Ht** |
| Number of measures length/lipids | 36 | 12/11 | 4 | 22 | 67/66 | 17 |
| Oil sac area, OSA, mm<sup>2</sup> | 0.03 ± 0.01 | 0.11 ± 0.01 | 0.00 ± 0.00 | 0.06 ± 0.01 | 0.43 ± 0.01 | 0.07 ± 0.02 |
| % WE max capacity | 5.2 | 19.4 | 0 | 5.6 | 66.3 | 7.3 |
| Respiration rate $\mu\text{gC ind}^{-1} \text{d}^{-1}$ | 0.08 | 0.1 | 0.09 | 0.2 | 0.27 | 0.21 |
| Development time (d) | 15 | 15 | 15 | 22 | 22 | 22 |
| Lipid accumulation $\mu\text{gC ind}^{-1} \text{d}^{-1}$ | 0.11 | 0.48 | 0.00 | 0.16 | 2.42 | 0.22 |

**Figure 1.**
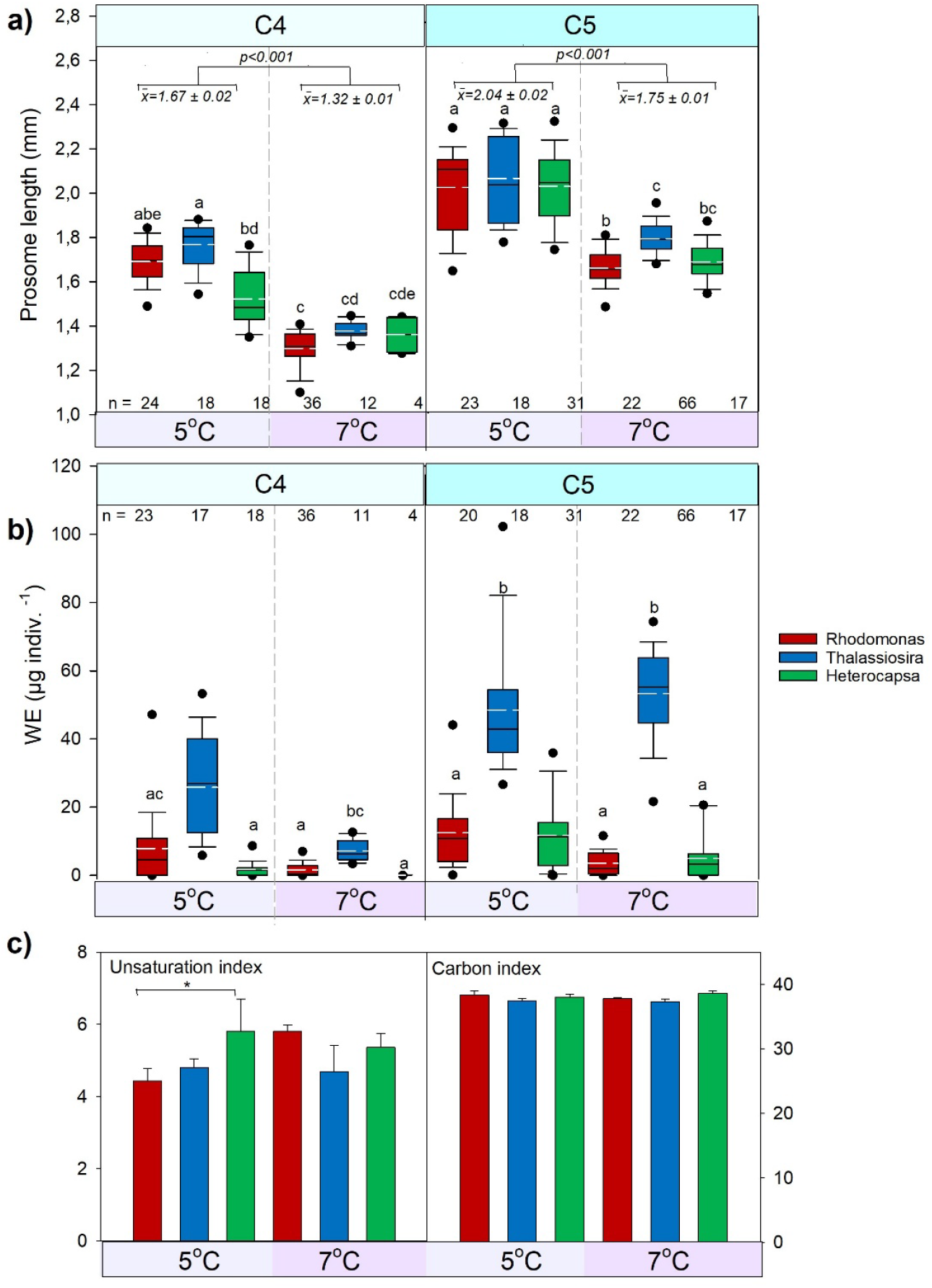
Calanus finmarchicus. Box diagrams of a) prosome length (mm), b) wax ester content (µg indiv^-1^) and c) unsaturation and carbon indexes (+1 stdev) of stage C4 and C5 WE after feeding the 3 different phytoplankton species *Rhodomonas salina* (red), *Thalassiosira weissflogii* (blue) and *Heterocapsa triquetra* (green) at 5° and 7°C. In a) and b) the box, whiskers, and solid dots outline 25/75, 90/10 and 95/5 percentiles. Solid line: median, dotted white line: mean. Lower case letters indicate no significant difference (P>0.05) between those sharing the same letter within stages n: number of measures. In a) average ± standard error of the prosome lengths and the p value of the ANOVA within stages, in c) * significant difference *p<0*.*05*.

**Figure 2.**
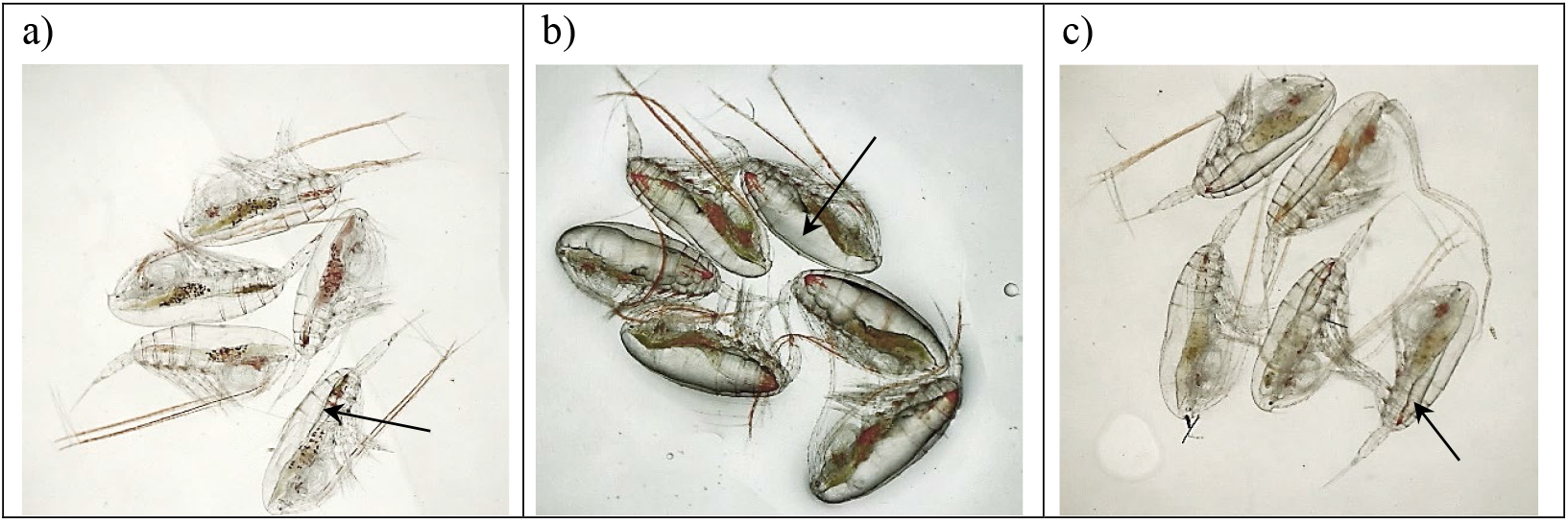
*Calanus finmarchicus* stage C5 at end of experiment at 7°C, after feeding on a) *R. salina*, b) *T. weissflogii* and c) *H. triquetra*. The oil sac is the clear structure shown by arrows.

Net lipid accumulation reflects the energy available after basic metabolism, so that observed differences in WE content could be a consequence of either different ingestion rates or assimilation efficiency. Regardless of the physiological reason, the diatom diet results in faster lipid accumulation, allowing earlier descent from the surface to deeper waters (12), reducing population predation pressure and increasing population survival. The copepods feeding on flagellate and dinoflagellate-based diets will on the other hand need considerably more time, longer than the bloom duration (240-331 days respectively) to reach a similar lipid quantity before entering diapause (Table 1).

These results indicate three compounding effects, each impacting the ability of *C. finmarchicus* to successfully overwinter. Firstly, lipid accumulation depends critically on the presence of diatoms being typically 3 times higher on a diatom than a flagellate-based diet. Secondly, increasing temperatures both in the surface and (potentially) at depth mean higher metabolic rates with less efficient accumulation and usage of lipids. Finally, reduced size of maturing individuals increases their specific respiration rates so that smaller individuals will require proportionally larger lipid reserves to fuel overwinter (7, 13).

While our focus is in the North Atlantic and Arctic, changing phytoplankton taxonomic composition is a common feature of many high latitude ecosystems (1, 2). It is evident that community structures are shifting away from diatoms and copepods towards a greater abundance of flagellates and gelatinous zooplankton (1). A unifying feature of these systems is the efficient transfer of lipids from diatoms to large zooplankton (copepods and krill) providing both reserves for overwintering and an efficient trophic link to higher trophic levels. Reduced ability of these large zooplankton to accumulate lipids poses an existential risk to these taxa, with repercussions for global productivity and carbon sequestration that remain poorly quantified.

## Methods

All data is presented in the manuscript, and detailed methodology is provided in supplementary information.

*Calanus finmarchicus* females were grown at two temperatures, 5 and 7 °C and fed mixture of the *Rs, Tw* and *Ht ad-libitum*. Eggs were hatched and the nauplii fed as above. After metamorphosis to copepodite stage C1, the population was split into three groups, each of which was fed pure diets of the experimental food at 400 μgC L^-1^ and let develop to stage C4 and C5. Individuals were photographed and measured for size and lipids. Copepods were extracted and their wax ester composition analyzed.

## Acknowledgments

This project was funded by the European Union’s Horizon research and innovation programme under grant agreements 869383 (ECOTIP) and 101136480 (SEA-Quester) and Research Council Faro Island 8011 (COPS). Torkel Nielsen provided copepod females from Greenland and Jack Melby and Markus Strange laboratory assistance.

## Supporting Information for

Diatom – lipid – copepod nexus under threat by global change

## Extended materials and methods

### Copepod culturing

*Calanus finmarchicus* females were collected in Disko Bay, Greenland in spring and transported live to Denmark. Females were cultured in a mixture of water from the Labrador Sea and artificial seawater at 33 psu. They were fed a mixture of the cryptophyte *Rhodomonas salina* (Rs), the diatom *Thalassiosira weissflogii* (Tw) and the dinoflagellate *Heterocapsa triquetra* (Ht) *ad libitum* (ESD: 7, 13 and 16µm respectively). The respective experimental temperatures of 5 and 7 °C were applied from the start of the female culturing.

Eggs were harvested and distributed into 20L beakers with filtered seawater at the experimental temperature. Approximately 1000 eggs were put in each beaker. When hatched, and nauplii were at the first feeding stage N2, they were fed the same food mixture as above at more controlled concentrations of 400 µgC L^-1^. After metamorphosis to C1, the population was split into three groups, each of which was fed pure diets of the experimental food, Rs, Tw and Ht at excess food concentration of 400 µgC L^-1^.

Naupliar (mixed diet) and copepodite (pure diet) cultures were maintained by changing 80% of the water 2-3 times a week, by reverse filtration of the culture vessel, replacing it with fresh seawater and food.

For all treatments, the experiment was terminated when copepodites reached copepodite stages 4 (C4) and 5 (C5) this took about 20-34 days for 7 and 5 depending on temperatures. All individuals were photographed through Leica-S9i stereomicroscope, and four to five individuals were sorted into dry tapered glass vials, topped with Nitrogen gas and frozen at -80°C.

From the images, the copepod prosome length and oil sac area (OSA) were traced using the “image-J” software (14). The pixel-to-mm ratio was calibrated using the image of the appropriate calibration slide. OSA was converted into wax esters (WE) using the formula WE (mg) = 0.167 × A^1.42^ from (15), where A is the area of the lipid sac in mm^2^.

### WE analysis

Copepods were weighed and counted into polypropylene tubes using 5-40 mg of sample and extracted in 10 µl of a 0.2 mg/ml butylated hydroxytoluene (BHT) solution in 50:50 water/methanol, followed by 350 µl of 90:10 methanol/water solution. Three glass beads were added to each vial and the sample homogenized using a bead beater at 2000 rpm for 30 seconds. Following this, 1000 µl of Methyl tert-butyl ether (MTBE) was added and the sample vortexed and left at room temperature for 5 minutes. Phase partition was ensured by adding 250 µl of water followed by vortex and centrifugation at 10,000 rpm for 5 minutes. The upper MTBE phase was then transferred with glass pipettes into clean Eppendorf tubes, and the original samples re-extracted in the same way with 1000 µl of MTBE which was subsequently pooled with the first extract. Samples were dried under a stream of nitrogen and resuspended in 1 ml of 2:1 MTBE/methanol solution. Depending on the amount of sample available, various dilutions ranging from 1 in 5 to 1 in 80 were used to allow for optimum column loading for each sample.

Samples were analyzed using Supercritical fluid chromatography (SFC-ESI-MS/MS) on an Acquity UPC2 system (Waters, USA) as previously described (16), using a method similar to the neutral lipid separation. Separation conditions were kept the same, with mass spectrum acquired over a smaller range, 50-1000 Da, using the data dependent mode and a ramped collision energy of 28-40 V for the product ion scan.

Unsaturation and carbon number indexes of the copepod WE were calculated representing the number of double bonds and carbon respectively.

Unsaturation index (average double bond number)

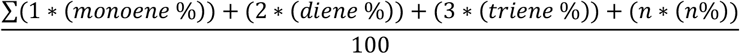

Average carbon number (wax-esters)

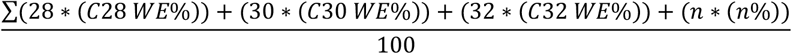

### Calculations in Table 1

WE µgC ind^-1^ calculated from OSA (15): WE (mgC) = 0.167 × A^1.42^, where A is the OSA mm^2^.

Maximum carbon-based WE content (WE_max_) is as described in (7) and is based on a structural carbon mass *m* scaled to prosome length (PL) as: *m* = αPL^3^ and WE carbon mass *w*_max_ = *β*PL^3^. The coefficients *α* and *β* are defined by a best-fit regression of maximum total carbon content per PL observed for different-sized copepods in their study (7).

Respiration rate is based on temperature (T) using (µmol O_2_ gC^-1^ hr^-1^ = 0.0441T + 1.1528) (13) and converted to carbon and adjusted to individual rates with copepod carbon conversion of (17).

Development time (DT) are the days taking to develop from stage C1 to C4 or C5 based on the Belehrádek’s function parameters for *C. finmarchicus*: DT= a(T+9.11)^-2.05^ where “a” is listed for each stage (18).

Lipid accumulation rate µgC ind^-1^ is attained by dividing carbon-based lipid content ind^-1^ with the development time

### Statistics

Difference between means was tested by one-way ANOVA with all pairwise multiple comparison by Holm-Sidak method. When normality test failed, we used Kruskal-Wallis One Way Analysis on ranks.

## References

1. S. A. Henson, B. B. Cael, S. R. Allen, S. Dutkiewicz, Future phytoplankton diversity in a changing climate. Nat Commun 12 (2021).

2. Hayward, A., et al., Antarctic phytoplankton communities restructure under shifting sea-ice regimes. Nature Climate Change (2025). 10.1038/s41558-025-02379-x.

3. M. Ardyna, K. R. Arrigo, Phytoplankton dynamics in a changing Arctic Ocean. Nat. Clim. Chang. 10, 892–903 (2020).

4. Z. Yi, M. Xu, X. Di, S. Brynjolfsson, W. Fu, Exploring Valuable Lipids in Diatoms. Front. Mar. Sci. 4 (2017).

5. I. Prokopchuk, E. Sentyabov, Diets of herring, mackerel, and blue whiting in the Norwegian Sea in relation to Calanus finmarchicus distribution and temperature conditions. ICES Journal of Marine Science 63, 117–127 (2006).

6. M. Frederiksen, M. Edwards, A. J. Richardson, N. C. Halliday, S. Wanless, From plankton to top predators: bottom-up control of a marine food web across four trophic levels. Journal of Animal Ecology 75, 1259–1268 (2006).

7. S.H. Jónasdóttir, R. J. Wilson, A. Gislason, M. R. Heath, Lipid content in overwintering Calanus finmarchicus across the Subpolar Eastern North Atlantic Ocean. Limnol Oceanogr 64, 2029–2043 (2019).

8. J. Pinti, S.H. Jónasdóttir, N. R. Record, A. W. Visser, The global contribution of seasonally migrating copepods to the biological carbon pump. Limnology and Oceanography 68, 1147–1160 (2023).

9. S.H. Jónasdóttir, A. W. Visser, K. Richardson, M. R. Heath, Seasonal copepod lipid pump promotes carbon sequestration in the deep North Atlantic. Proc Natl Acad Sci USA 112, 12122–12126 (2015).

10. A. W. Visser, S.H. Jónasdóttir, Lipids, buoyancy and the seasonal vertical migration of Calanus finmarchicus. Fisheries Oceanography 8, 100–106 (1999).

11. R. J. Wilson, N. S. Banas, M. R. Heath, D. C. Speirs, Projected impacts of 21st century climate change on diapause in Calanus finmarchicus. Global change biology 22, 3332–3340 (2016).

12. M. S. Schmid, F. Maps, L. Fortier, Lipid load triggers migration to diapause in Arctic Calanus copepods—insights from underwater imaging. Journal of Plankton Research 40, 311–325 (2018).

13. W. J. Saumweber, E. G. Durbin, Estimating potential diapause duration in Calanus finmarchicus. Deep Sea Research Part II: Topical Studies in Oceanography 53, 2597–2617 (2006).

## SI References

1. W. S. Rasband, National Institutes of Health, Bethesda, Maryland, USA. http://imagej.nih.gov/ij/ (2011).

2. D. Vogedes, et al., Lipid sac area as a proxy for individual lipid content of arctic calanoid copepods. Journal of Plankton Research 32, 1471–1477 (2010).

3. R. Broughton, D. R. Tocher, M. B. Betancor, Development of a C18 Supercritical Fluid Chromatography-Tandem Mass Spectrometry Methodology for the Analysis of Very-Long-Chain Polyunsaturated Fatty Acid Lipid Matrices and Its Application to Fish Oil Substitutes Derived from Genetically Modified Oilseeds in the Aquaculture Sector. ACS Omega 5, 22289–22298 (2020).

4. S.H. Jónasdóttir, R. J. Wilson, A. Gislason, M. R. Heath, Lipid content in overwintering Calanus finmarchicus across the Subpolar Eastern North Atlantic Ocean. Limnol Oceanogr 64, 2029–2043 (2019).

5. W. J. Saumweber, E. G. Durbin, Estimating potential diapause duration in Calanus finmarchicus. Deep Sea Research Part II: Topical Studies in Oceanography 53, 2597–2617 (2006).

6. A. Ingvarsdóttir, D. F. Houlihan, M. R. Heath, S. J. Hay, Seasonal changes in respiration rates of copepodite stage V Calanus finmarchicus (Gunnerus). Fisheries Oceanography 8, 73–83 (1999).

7. R. G. Campbell, M. M. Wagner, G. J. Teegarden, C. A. Boudreau, E. G. Durbin, Growth and development rates of the copepod Calanus finmarchicus reared in the laboratory. Marine Ecology Progress Series 221, 161–183 (2001).

